# Phase Resolution of Heterozygous Sites in Diploid Genomes is Important to Phylogenomic Analysis under the Multispecies Coalescent Model

**DOI:** 10.1101/2021.03.29.437575

**Authors:** Jun Huang, Jeremy Bennett, Tomáš Flouri, Adam D. Leaché, Ziheng Yang

## Abstract

Genome sequencing projects routinely generate haploid consensus sequences from diploid genomes, which are effectively chimeric sequences with the phase at heterozygous sites resolved at random. The impact of phasing errors on phylogenomic analyses under the multispecies coalescent (MSC) model is largely unknown. Here we conduct a computer simulation to evaluate the performance of four phase-resolution strategies (the true phase resolution, the diploid analytical integration algorithm which averages over all phase resolutions, computational phase resolution using the program PHASE, and random resolution) on estimation of the species tree and evolutionary parameters in analysis of multi-locus genomic data under the MSC model. We found that species tree estimation is robust to phasing errors when species divergences were much older than average coalescent times but may be affected by phasing errors when the species tree is shallow. Estimation of parameters under the MSC model with and without introgression is affected by phasing errors. In particular, random phase resolution causes serious overestimation of population sizes for modern species and biased estimation of cross-species introgression probability. In general the impact of phasing errors is greater when the mutation rate is higher, the data include more samples per species, and the species tree is shallower with recent divergences. Use of phased sequences inferred by the PHASE program produced small biases in parameter estimates. We analyze two real datasets, one of East Asian brown frogs and another of Rocky Mountains chipmunks, to demonstrate that heterozygote phase-resolution strategies have similar impacts on practical data analyses. We suggest that genome sequencing projects should produce unphased diploid genotype sequences if fully phased data are too challenging to generate, and avoid haploid consensus sequences, which have heterozygous sites phased at random. In case the analytical integration algorithm is computationally unfeasible, computational phasing prior to population genomic analyses is an acceptable alternative.

## 1. Introduction

Next-generation sequencing technologies have revolutionized population genetics and phylogenetics by making it affordable to sequence whole genomes or large portions of the genome, even for non-model organisms. Many phylogenomic studies use the approach of reduced representation library to maximize their DNA sequencing efforts on a small subset of the genome. These strategies can generate thousands of genomic segments (called loci in this paper irrespective of whether they are protein-coding) with high coverage, and target sequences can be assembled with confidence. Examples include restriction site-associated DNA sequencing (RADseq), which is used frequently to identify single nucleotide polymorphisms (SNPs) for population genetic and phylogeographic studies (Andrews *et al*., 2016; Leaché and Oaks, 2017), although it has also been applied to address phylogenetic questions at deeper timescales (Eaton *et al*., 2017). A more common approach for phylogenomic studies is targeted sequence capture, generating so-called reduced-representation datasets, with typically longer sequences for distantly related species than with RADseq data. Examples include exome sequencing, ultraconserved elements (UCEs, Faircloth *et al*., 2012), anchored hybrid enrichment (AHE, Lemmon *et al*., 2012), conserved nonexonic elements (CNEEs, Edwards *et al*., 2017), or rapidly evolving long exon capture (RELEC, Karin *et al*., 2020).

Typical sequencing technologies produce short fragments of sequenced DNA called ‘reads’ that are either *de novo* assembled or mapped to a pre-existing reference genome. This leads to chromosomal positions being sequenced a variable number of times across the genome (usually referred to as the sequencing depth). A common practice in genome sequencing projects has been to produce the so-called “haploid consensus sequence” for a diploid individual, which uses the most common nucleotide at any heterozygous site to produce one genomic sequence. Assemblers like Velvet (Zerbino and Birney, 2008), ABySS (Simpson *et al*., 2009), and Trinity (Grabherr *et al*., 2011), pick up only one of the two nucleotide bases at any heterozygous site and essentially resolve the phase of heterozygous sites at random, producing chimeric sequence that may not exist in nature. Suppose a diploid individual is heterozygous at two sites in a genomic region, so that the diploid genotype may be represented Y…R, with two heterozygous sites Y (for T/C) and R (for A/G) (Fig. 1). Suppose the reads are 14 *×* T and 6 *×* C at the first site, and 7 *×* A, 10 *×* G, and 1 *×* T at the second (with the single T to be most likely a sequencing error). The haploid consensus sequence is constructed as T…G. In effect a heterozygote site with high quality scores for the two nucleotides is represented as one consensus nucleotide with a low quality score. Because it is largely pure chance which of the two nucleotides at a heterozygous site has the greater number of reads, this strategy is equivalent to resolving the phase at random and using only one of the constructed sequences. The resulting haploid consensus sequence may not be a real biological sequence and may not represent the biology of the diploid individual. Besides loss of information, a more serious problem is that the artefactual phased haploid sequence may be unusually divergent from other sequences in the sample, potentially introducing systematic biases in downstream inference. Currently constructing true diploid *de novo* assemblies is expensive. A sequencing platform has been developed in combination with bioinformatic algorithms to determine the true diploid genome sequence but the strategy still involves high cost (Weisenfeld *et al*., 2017). If a read is long and fully covers a locus, multiple heterozygous sites in the same locus will be naturally phased. However, if the reads are short, and the two heterozygous sites do not occur in the same read, their genotypic phase resolution will become an issue.

**Fig. 1.**
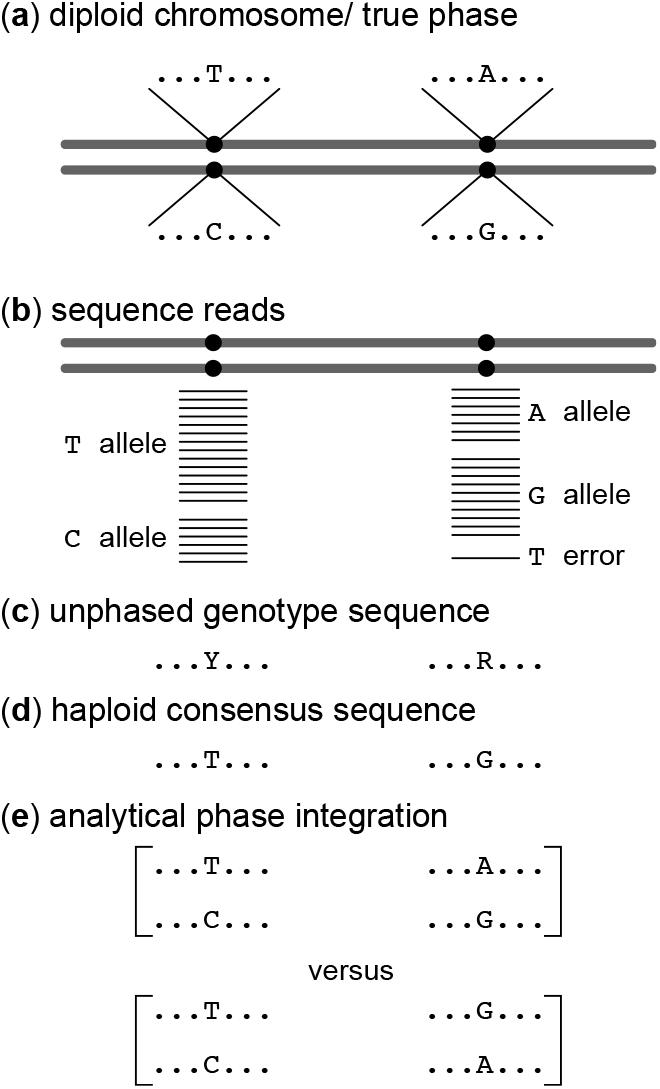
Example of heterozygote phase resolution. (a) A hypothetical diploid chromosome with two heterozygous sites (T/C and A/G). The true haploid genotypes are T…A and C. G. (b) Sequence reads around the two heterozygous sites, assuming that they are far apart on the chromosome so that they are not present on any single read (in which case phase would be determined) while they are close enough to be on one locus. In this case genome assemblers should produce the unphased genotype sequence (c), using the IUPAC ambiguity codes to represent heterozygote sites, but instead they produce the so-called ‘haploid consensus sequence’ (d), picking up the most common nucleotide at each heterozygote site (T G since T and G are by chance the most common sequence reads at the two sites), which may not match either of the true haploid sequences. (e) Analytical integration of phase resolution takes the unphased genotype sequences as data and averages over all possible phase resolutions, weighting each one appropriately according to their relative likelihood based on the whole sequence alignment at the locus.

How the heterozygote phase is resolved may have a significant impact on population genomic and phylogenomic inference using genomic sequence data. Phase information is well-known to be important for relating genotype to phenotype in human disease mapping (Tewhey *et al*., 2011). Similarly, Gronau *et al*. (2011) found that use of an analytical integration method (which averages over all possible phase resolutions) leads to nearly identical performance as the use of true phase resolutions for estimating population parameters, and that random phase resolution produced unreliable estimates. Andermann *et al*. (2019) developed a bioinformatics pipeline to recover allelic sequences from sequence capture data, and found it to produce more accurate estimation of species divergence times under the MSC model (Rannala and Yang, 2003) than other strategies such as use of consensus haploid sequences, random phasing, or ambiguity encoding. Overall little is known about the effects of heterozygote phase resolution on many inference problems using multilocus genomic sequence data under the MSC model, including species tree estimation, estimation of population sizes and species divergence times, and inference of cross-species introgression/hybridization.

We have implemented in BPP (Flouri *et al*., 2018) an analytical integration algorithm to handle unphased diploid sequences, developed by Gronau *et al*. (2011) in their G-PhoCS program, which is an orthogonal extension of an earlier version of BPP (Rannala and Yang, 2003; Burgess and Yang, 2008). Previously Kuhner and Felsenstein (2000) implemented an Markov chain Monte Carlo (MCMC) algorithm to average over different phase resolutions in the likelihood calculation for estimating *θ* under the single-population coalescent. The algorithm was found to mix slowly even for small datasets. The analytical integration algorithm uses a data-augmentation strategy, in which the unknown fully resolved haploid sequences constitute the complete data or latent variables, and enumerates and averages over all possible phase resolutions, weighting them according to their likelihoods based on the whole sequence alignment. For example, if a diploid sequence has two heterozygous sites, Y…R, the approach will average over both phased genotypic resolutions: (i) T…A and C…G versus (ii) T…G and C…A (Fig. 1). Note that there may be rich information about the phase resolution of any unphased sequence in an alignment of many sequences, either from the same species or from different but closely related species. Consider for example the phase resolutions for a human diploid sequence Y…R (Fig. 1). If we observe in the chimpanzee fully resolved sequences T…A and C…G (e.g., in an individual homozygous at both sites, with genotypes T/T…A/A) and never observe sequences T…G and C…A, then very likely the human diploid sequence has the haploid genotypes T…A and C…G. Our implementation of the algorithm works with all four analyses under the MSC model in BPP (Yang, 2015; Flouri *et al*., 2018, 2020b), including species tree estimation (Yang and Rannala, 2014; Rannala and Yang, 2017) and species delimitation through Bayesian model selection (Yang and Rannala, 2010, 2014; Leaché *et al*., 2019). We also implemented the algorithm under the multispecies-coalescent-with-introgression (MSci) model (Flouri *et al*., 2020a).

Here we use computer simulation to evaluate different phase-resolution strategies in terms of their precision and accuracy in Bayesian species tree estimation under the MSC and in parameter estimation under both the MSC and MSci models. In addition to using the true phase resolution, which is generated during the simulation and is known with certainty, we also include analytical phase integration (Gronau *et al*., 2011; Flouri *et al*., 2018), phase resolution using the program PHASE (Stephens *et al*., 2001; Stephens and Donnelly, 2003), and random resolution. The strategy of random resolution is largely equivalent to the common method of using haploid consensus sequences. The PHASE program was developed for population data from the same species, but is here applied to unphased sequences from both within and between species. We note that a number of computational phasing algorithms have been developed, such as Haplotyper (Niu *et al*., 2002) and fastPHASE (Scheet and Stephens, 2006). These are mostly developed to improve the computational efficiency and to handle long sequences (Choi *et al*., 2018), and are expected to produce similar results to PHASE in analysis of short sequences.

## 2. Methods andMaterials

### Simulation to Estimate Species Trees

We use the program MCcoal in BPP3.4 (Yang, 2015) or the simulate switch of BPP4.3 (Flouri *et al*., 2020b) to simulate gene trees and multi-locus sequence data using four fixed species trees for eight species (Figs. 2a, a′, b, & b′). The trees have very short branches, mimicking challenging species trees generated during radiative speciation events. In the two deep trees species divergences are much older than average coalescent times (*θ/*2). In the two shallow trees, species divergences are very recent relative to coalescent times, mimicking different populations of the same species. Note that in this study, we make no distinction between species and populations. The MSC model has two sets of parameters: the species divergence times (*τ*s) and the population size parameters (*θ*s). Both are measured by the expected number of mutations/substitutions per site. For each species/population, *θ* = 4*Nµ*, where *N* is the effective population size and *µ* is the mutation rate per site per generation. We consider two mutation rates, with *θ* = 0.001 (low rate) or 0.01 (high rate), respectively, for all populations on the tree. The species divergence times (*τ*s) are given as multiples of *θ* . We consider 10, 20, 50, or 100 loci, with each locus having 500 sites. On average there should be 0.5 and 5 heterozygous sites between the two sequences of any individual at the low and high rates, respectively. We sample *S* = 2 or 4 haploid sequences (or 1 or 2 diploid individuals) per species at each locus. Gene trees with branch lengths (coalescent times) are generated independently among loci using the MSC density given the species tree and parameters (Rannala and Yang, 2003). The JC model (Jukes and Cantor, 1969) is then used to ‘evolve’ the sequences along the gene tree to generate the sequence alignments at the tips of the tree. Analysis of this full dataset by BPP is strategy ‘F’.

**Fig. 2.**
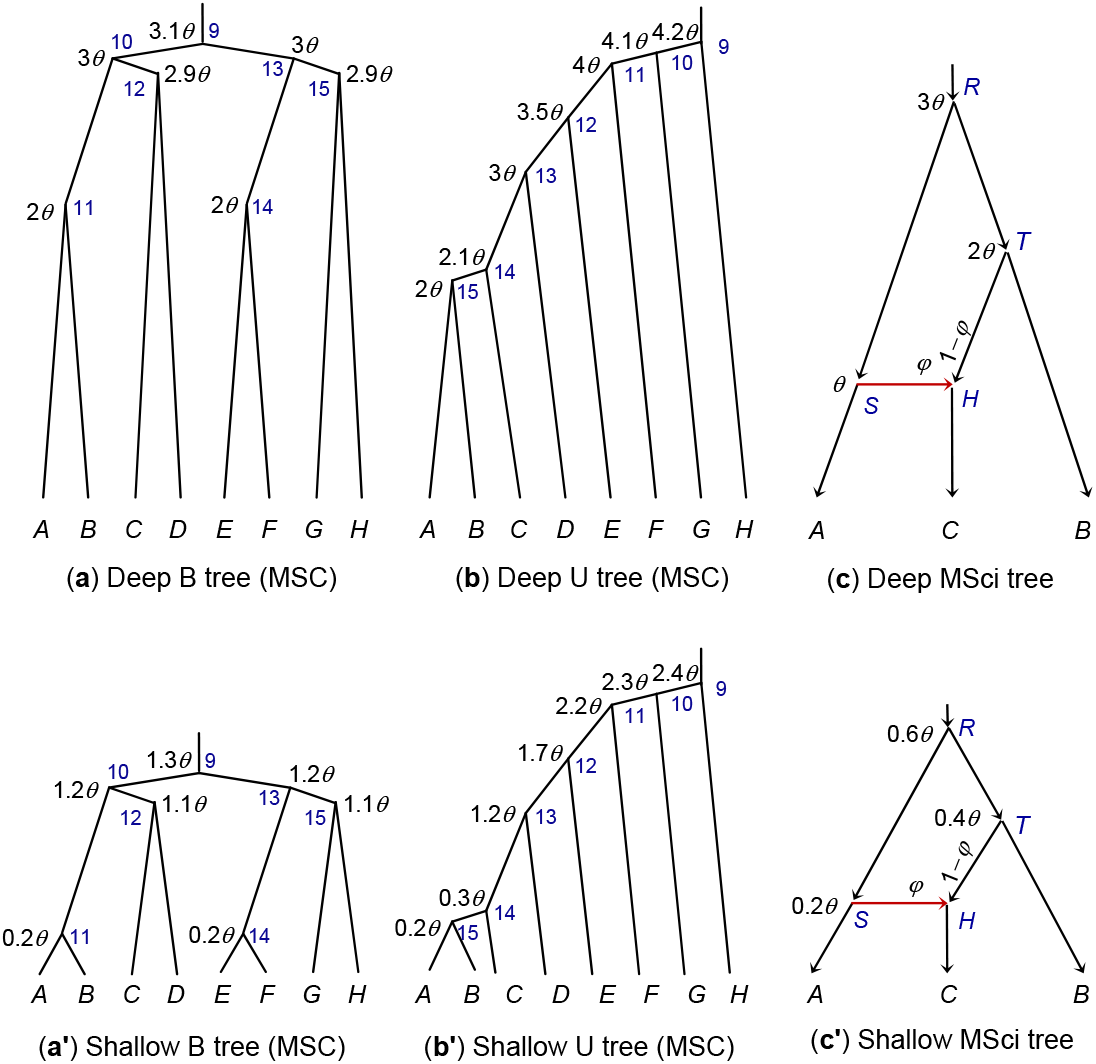
(**a & a**′) Deep and shallow balanced species trees, and (**b & b**′) deep and shallow unbalanced species trees for eight species used for simulating data under the MSC model. (**c & c**′) Deep and shallow species trees with introgression used to simulate data under the MSci model. The ages of internal nodes (*τ*s) are shown next to the nodes, with *θ* = 0.01 (high rate) or 0.001 (low rate). The blue indexes at internal nodes of the tree are used to identify the parameters.

To simulate unphased diploid sequences, two sequences from the same species are combined into one diploid sequence, using the International Union of Pure and Applied Chemistry (IUPAC) ambiguity characters to represent heterozygous sites (for example, Y means a T/C heterozygote) (Fig. 1c). The data of unphased diploid sequences are analyzed using the diploid or phase option of the BPP program (strategy ‘D’), which analytically averages over all possible phase resolutions (Gronau *et al*., 2011). With strategy ‘P’, we use the program PHASE (Stephens *et al*., 2001) to resolve the phase, and then analyze the phased sequences using BPP (with 16 or 32 sequences in the alignment per locus for *S* = 2 and 4, respectively). Lastly, we use random phase resolution, referred to as strategy ‘R’. The simulation program automatically generates the sequence alignments for strategies F, D, and R. For strategy P, we ran PHASE 2.1 (Stephens *et al*., 2001) to reconstruct the phased sequences for each locus, and used the PERL program SeqPhase (Flot, 2010) to convert files.

The number of replicate datasets is 100. With four trees, two mutation rates (*θ* = 0.001 or 0.01), two sampling configurations (*S* = 2 or 4), four numbers of loci (*L* = 10, 20, 50, 100), we generated in total 4 *×* 2 *×* 2 *×* 4 *×* 100 = 6400 datasets, each of which is analyzed using the four strategies. The BPP program (Flouri *et al*., 2018) was used in the analysis. Inverse-gamma priors are assigned on parameters under the MSC model, with the shape parameter 3 so that the priors are diffuse and with the mean to be close to the true value. We use *θ∼* IG(3, 0.02) with mean 0.01 and *τ*_0_ *∼* IG(3, 0.08) with mean 0.04 for the age of the root of the species tree for data simulated with the high rate (*θ* = 0.01). For data of the low rate (*θ* = 0.001), the priors are *θ ∼* IG(3, 0.002) with mean 0.001 and *τ*_0_ *∼* IG(3, 0.008) with mean 0.004. The prior means for *τ*_0_ are close to the true values for the deep trees but are larger than the true values for the shallow trees, although the priors are diffuse. For species tree estimation, we integrate out *θ*s analytically through the use of the conjugate inverse-gamma priors. We conducted pilot runs to determine the chain lengths needed for convergence. The final settings for the MCMC are 20,000 iterations for burn-in, then taking 2 *×* 10^5^samples, sampling every 2 iterations.

Strategy P requires running the Bayesian MCMC program PHASE *L* times if there are *L* loci in the dataset, to generate the fully resolved sequence alignments at the loci. This is somewhat expensive if there is a large number of loci and the mutation rate is high resulting in many heterozygous sites at each locus. After the datasets are generated, the BPP analysis of each dataset by strategies F, P, and R involves about the same amount of computation. Strategy D is more expensive as the method averages over all possible phase resolutions, which may involve likelihood calculation for many site patterns, especially if there are many sequences per locus with many heterozygous sites.

For species tree estimation (A01 analysis in Yang, 2015), we calculated the proportion (among the 100 replicates) with which each node on the true species tree is found in the *maximum a posteriori* (MAP) species tree in the BPP analysis. This is a measure of accuracy since the MAP tree is the best ‘point estimate’ of the species tree (Rannala and Yang, 1996). We examined the size and coverage probability of the 95% credibility set of species trees. The coverage probability is the proportion among the 100 replicate datasets in which the credibility set includes the true species tree. The size of the set indicates the precision or power of the method, but the method is considered reliable only if the coverage probability exceeds the nominal 95%.

### Simulation to Estimate Parameters under the MSC Model

The same data simulated under the MSC model for species tree estimation are analyzed using the four phase-resolution strategies to estimate parameters in the MSC model (*θ*s and *τ*s), with the species tree fixed. This is the A00 analysis in Yang (2015). We calculated the posterior means and the 95% HPD CI intervals for each parameter and examine the relative root mean square error (rRMSE), using the posterior means as point estimates. This is defined as

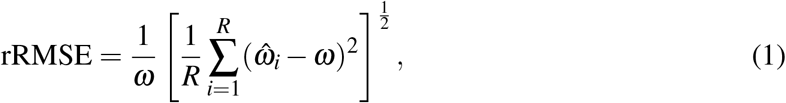

where *ω* is the true value of any parameter, and 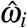 its estimate (posterior mean) in the *i*th replicate dataset, with *i* = 1, *…, R* over the *R* = 100 replicates. For example, rRMSE = 0.1 means that the mean square error is 10% of the true value. The rRMSE is a combined measure of bias and variance.

### Simulation to Estimate Parameters under the MSci Model

The MSci models for three species of Figures 2c&c′ are assumed to generate gene trees and sequence alignments using the simulate option of BPP4.3 (Flouri *et al*., 2020a). The three species have the phylogeny (*A*, (*C, B*)), but there was introgression from *A* to *C* at the time *τ*_*H*_ = *τ*_*S*_, with the introgression probability *ϕ* = 0.1 and 0.3. Other settings are the same as above for the simulation under the MSC model. We consider two mutation rates (with *θ* = 0.001 and 0.01) and four datasizes (with *L* = 10, 20, 50, and 100 loci), with each locus having 500 sites. We sample either *S* = 2 or 4 sequences per species per locus. The JC model is used both to simulate and to analyze the data.

For data simulated at the high rate (*θ* = 0.01), the priors are *θ∼* IG(3, 0.02) and *τ*_0_ *∼* IG(3, 0.06) for the root age. At the low rate (*θ* = 0.001), the priors are *θ ∼* IG(3, 0.002) and *τ*_0_ IG(3, 0.006). A 𝕌(0, 1) prior is used for the introgression probability *ϕ*.

### Analyses of two real datasets

We applied different phase-resolution strategies (D, P, and R) to analyze two previously published datasets, one of East Asian brown frogs (Zhou *et al*., 2012) and another of Rocky Mountains chipmunks (Sarver *et al*., 2021), to demonstrate that the effects discovered in the simulations apply to real data analysis. With real data, the option of true phase resolution (F) is unavailable, and the analytical phase resolution (D) is expected to perform the best. In addition, we include an approach of treating heterozygote sites in the alignment as ambiguity characters in the likelihood calculation, and refer to it as strategy ‘A’ (for ambiguity). This is considered a mistaken approach of handling the data and is not included in our simulation, but we use it in the real data analysis to illustrate its effects.

We re-analyzed a dataset of five nuclear loci from the East Asia brown frogs in the *Rana chensinensis* species complex (Zhou *et al*., 2012) to infer the species tree (the A01 analysis) and to estimate the parameters under the MSC on the MAP tree (the A00 analysis). There are three morphologically recognized species or four populations: *R. chensinensis* (clades C and L), *R. kukunoris* (K) and *R. huanrensis* (H) (Fig. 3a). The dataset was previously analyzed by Yang (2015), treating heterozygotes as ambiguities (strategy A). Each locus has 20-30 sequences, with sequence lengths to be 285–498 sites. We assign inverse-gamma priors on parameters: *θ ∼* IG(3, 0.002) with mean 0.001 and *τ*_0_ *∼* IG(3, 0.004) with mean 0.002 for the root age. We used a burnin of 8000 iterations, then taking 10^5^ samples, sampling every two iterations. The same analysis was run at least twice to confirm consistency between runs. This is a small dataset and the MCMC algorithm mixes well.

**Fig. 3.**
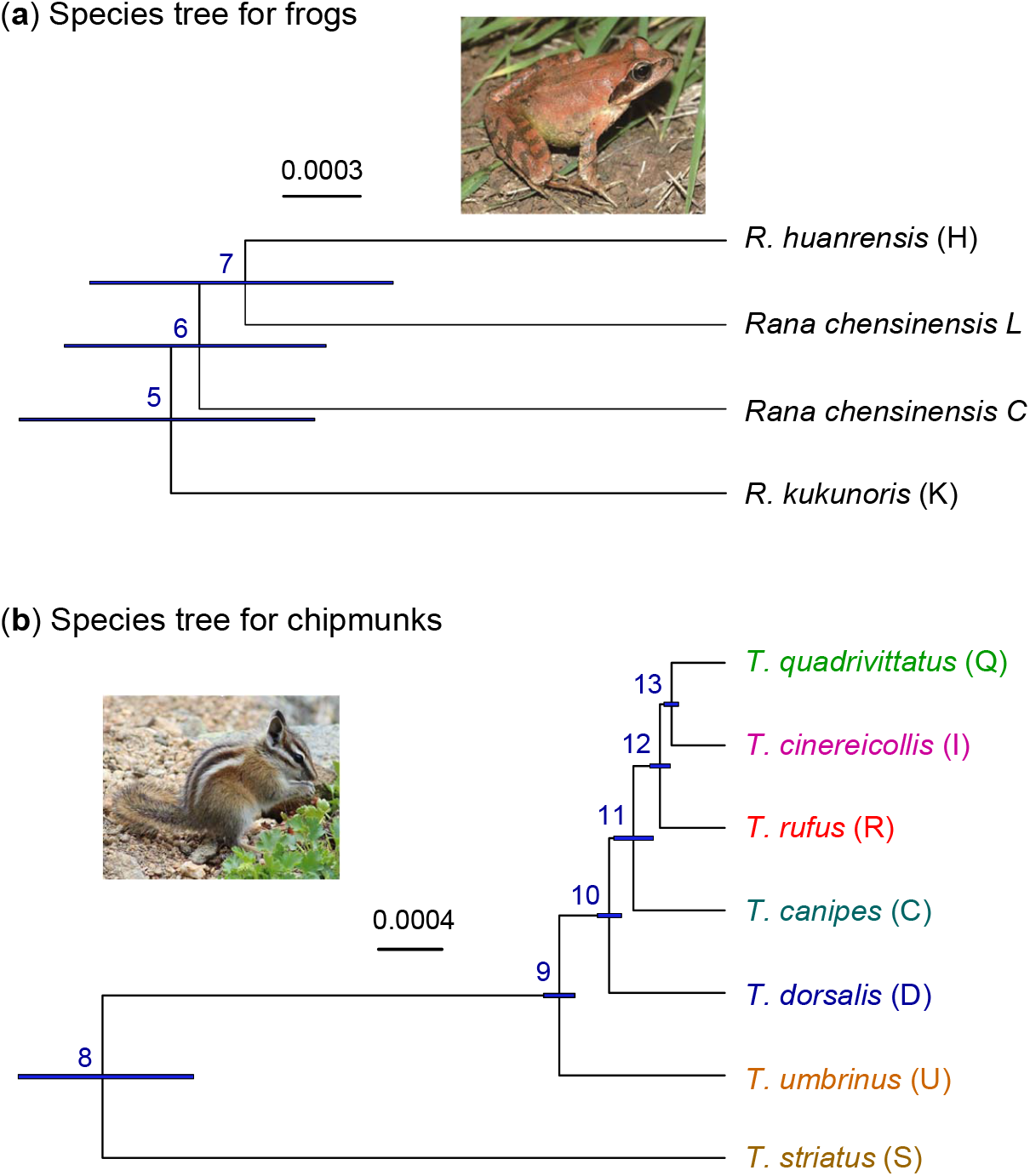
Inferred species trees (**a**) for East Asian brown frogs and (**b**) for Rocky Mountains chipmunks. Branch lengths reflect the posterior means of divergence times, with branch bars representing the 95% HPD intervals, obtained under the MSC using the analytical phase integration algorithm (strategy D). Estimates of other parameters are in table 6.

The second dataset consist of nuclear loci from six species of Rocky Mountains chipmunks in the *Tamias quadrivittatus* group: *T. canipes* (C), *T. cinereicollis* (I), *T. dorsalis* (D), *T. quadrivittatus* (Q), *T. rufus* (R), and *T. umbrinus* (U) (Fig. 3b). Sarver *et al*. (2021) used a targeted sequence-capture approach to sequence 51 Rocky Mountains chipmunks from those six species. As a reference genome assembly was lacking, reads were assembled iteratively into contigs using an approach called “assembly by reduced complexity”. A dataset of 1060 nuclear loci was compiled for molecular phylogenomic and introgression analyses, including 3 individuals from an outgroup species, *T. striatus*. High-quality heterozygotes, judged by mapping quality and read depth, are represented in the alignments using the IUPAC ambiguity codes. The filters applied by the authors suggest that the loci may be mostly coding exons or conserved parts of the genome. The majority of loci have 5 ≤ variable sites (including the outgroup). We used the first 500 loci in our analyses to infer the species tree and to estimate parameters under the MSC model. We assigned inverse-gamma priors on parameters: *θ ∼* IG(3, 0.002) with mean 0.001 and *τ*_0_ *∼* IG(3, 0.01) with mean 0.005 for the root age. In the A01 analysis (species tree estimation), we used a burnin of 16000 iterations, then taking 2 *×* 10^5^ samples, sampling every two iterations. The A00 analysis (parameter estimation on the MAP tree) used the same settings except that only 10^5^ samples were collected. The same analysis was run at least twice to confirm consistency between runs.

## 3. Results

### Species Tree Estimation under the MSC Model

Bayesian analysis of each replicate dataset using each of the four strategies produced a sample from the posterior distribution of the species trees, which we summarized to identify the maximum *a posteriori* probability (MAP) tree, and construct the 95% credibility set of species trees. The proportion, among the 100 replicates, with which the clades represented by those short branches were recovered in the MAP tree are shown in tables 1, S1–S3. Other clades on the trees, represented by longer branches, were recovered with probability near 100%, even for the low mutation rate and 10 loci. We also plotted the posterior probabilities for the true tree for the different phasing strategies in Figures 4, S1–S3. Strategy F, the analysis of the fully resolved haploid data, is expected to have the best performance and is thus the gold standard, against which the other strategies are compared.

**Table 1.**
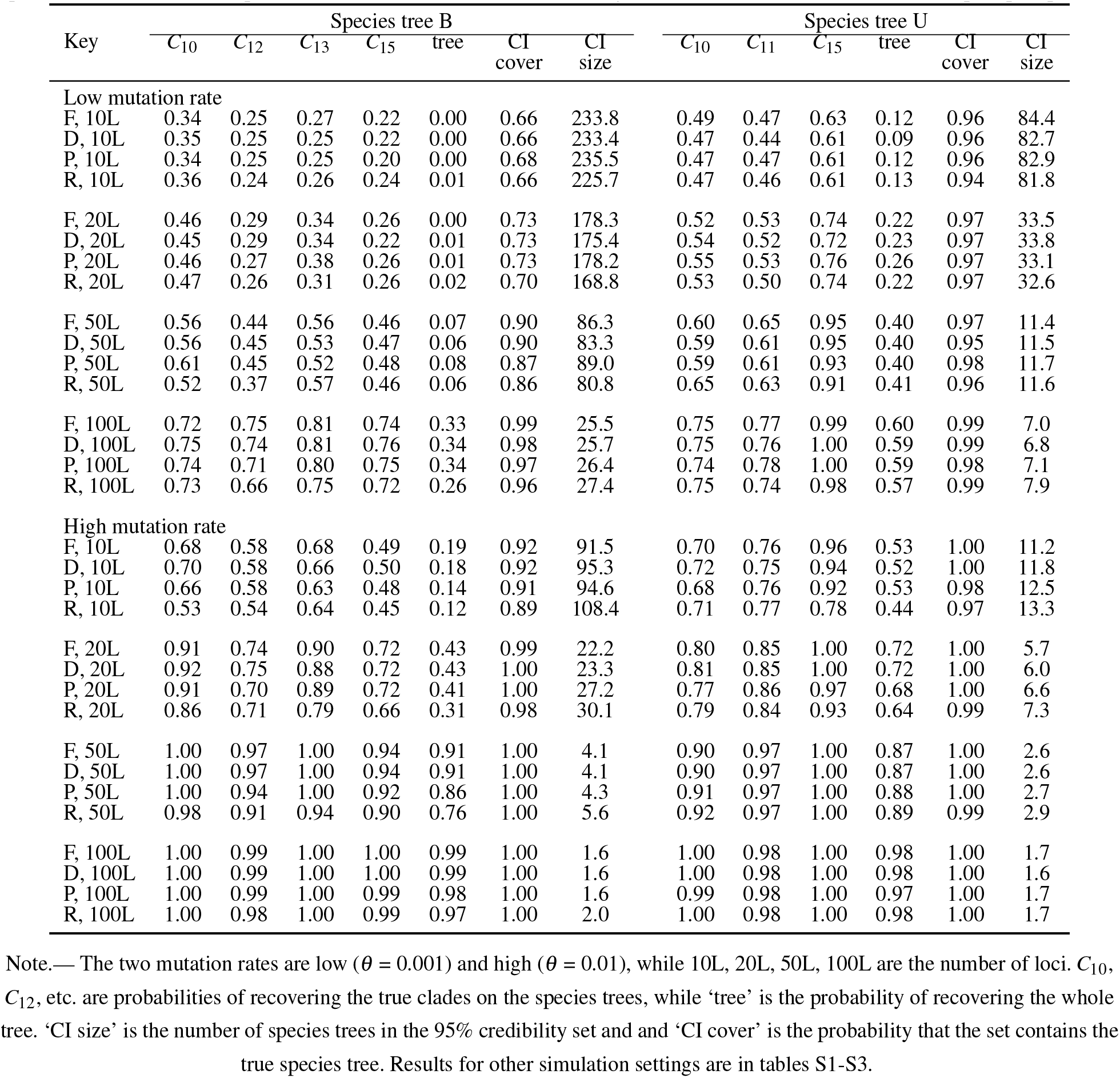
(**MSC A01, shallow**, *S* = 4) Probabilities of recovering true clades and the size and coverage of the 95% credibility set of species trees when the true species tree is Shallow B and Shallow U (Figs. 2a′&b′) and *S* = 4 sequences are sampled per species

**Fig. 4.**
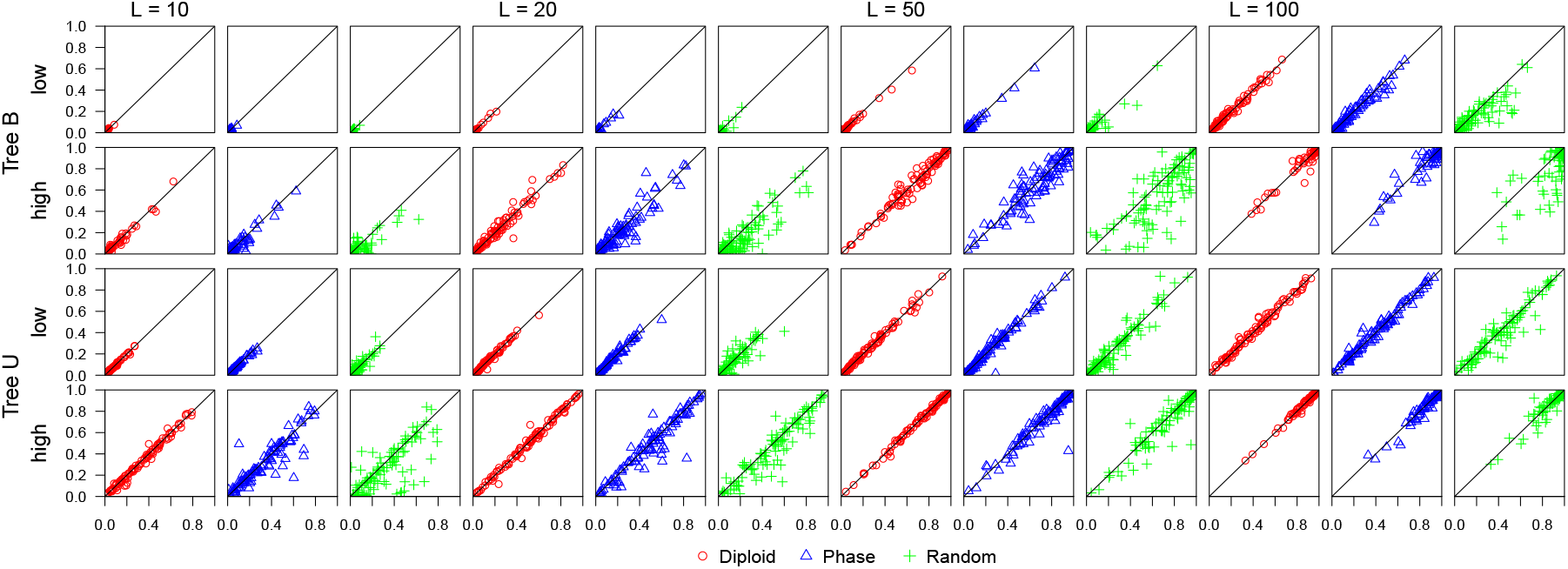
(**A01 under MSC, shallow tree**, *S* = 4) Posterior probability for the true species tree for phase-resolution strategies D (diploid), P (PHASE) and R (random) plotted against the probability for strategy F (full data). The data are simulated under the MSC models with species trees Shallow B and Shallow U (Figs. 2a′&b′), with *S* = 4 sequences sampled per species. Each plot has 100 scatter points, for the 100 replicate datasets, with the *x*-axis to be the posterior probability for strategy F while the *y*-axis is for strategies D, P, or R. ‘Low’ (*θ* = 0.001) and ‘high’ (*θ* = 0.01) refer to the mutation rate, and *L* (= 10, 20, 50, 100) is the number of loci. Results for other simulation settings are in Figures S1-S3.

In data simulated using the two deep trees (Deep B and Deep U) (Figs. 2a&b), the four phase-resolution strategies produced similar probabilities for recovering the true clades, with the differences among methods not being larger than the random sampling errors due to the limited number of replicates (tables S1 & S3). The different strategies most often produced the same MAP tree, although the posterior probability attached to the MAP tree varies somewhat among methods, but the differences are comparable to MCMC sampling errors. This can be seen in Figures S1 & S3, where the posterior for the true tree is plotted. Even random resolution (R) produced very similar results to the use of the fully resolved data (F). Note that in data simulated at the high rate, there are very likely to be two or more heterozygote sites in the diploid genotype of each individual at any locus, and the switching error rate for random phase resolution, which is the average proportion of heterozygous sites mis-assigned relative to the previous heterozygous site (Stephens and Donnelly, 2003), is 50%. Even the PHASE program generates substantial errors of phase resolution at the high mutation rate (table 2). Species tree estimation is thus robust to considerable phasing errors when species divergences are much older than average coalescent times.

**Table 2.**
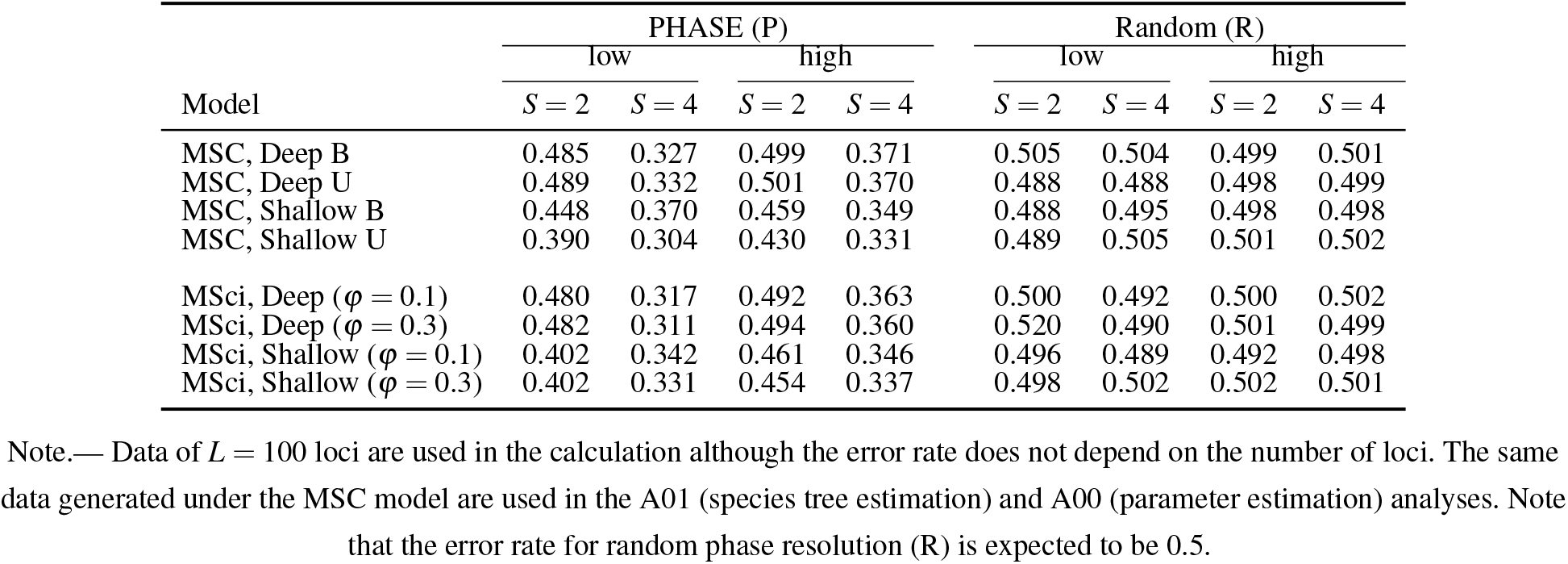
Average switching error rate for datasets simulated under the MSC and MSci models in this study

| Model | PHASE (P) |  |  |  | Random (R) |  |  |  |
| --- | --- | --- | --- | --- | --- | --- | --- | --- |
|  | low |  | high |  | low |  | high |  |
| | $S = 2$ | $S = 4$ | $S = 2$ | $S = 4$ | $S = 2$ | $S = 4$ | $S = 2$ | $S = 4$ |
| MSC, Deep B | 0.485 | 0.327 | 0.499 | 0.371 | 0.505 | 0.504 | 0.499 | 0.501 |
| MSC, Deep U | 0.489 | 0.332 | 0.501 | 0.370 | 0.488 | 0.488 | 0.498 | 0.499 |
| MSC, Shallow B | 0.448 | 0.370 | 0.459 | 0.349 | 0.488 | 0.495 | 0.498 | 0.498 |
| MSC, Shallow U | 0.390 | 0.304 | 0.430 | 0.331 | 0.489 | 0.505 | 0.501 | 0.502 |
| MSci, Deep ($\phi = 0.1$) | 0.480 | 0.317 | 0.492 | 0.363 | 0.500 | 0.492 | 0.500 | 0.502 |
| MSci, Deep ($\phi = 0.3$) | 0.482 | 0.311 | 0.494 | 0.360 | 0.520 | 0.490 | 0.501 | 0.499 |
| MSci, Shallow ($\phi = 0.1$) | 0.402 | 0.342 | 0.461 | 0.346 | 0.496 | 0.489 | 0.492 | 0.498 |
| MSci, Shallow ($\phi = 0.3$) | 0.402 | 0.331 | 0.454 | 0.337 | 0.498 | 0.502 | 0.502 | 0.501 |

Note.— Data of $L = 100$ loci are used in the calculation although the error rate does not depend on the number of loci. The same data generated under the MSC model are used in the A01 (species tree estimation) and A00 (parameter estimation) analyses. Note that the error rate for random phase resolution (R) is expected to be 0.5.

For the two shallow trees (Figs. 2a′&b′), large differences were found among the four strategies (tables 1 & S2, Figures 4 & S2). While strategy D produced results very similar to use of the full data (F), both strategies P and R had poorer performance, especially at the high rate, when strategy R produced larger CI set, with lower coverage than strategies F and D.

Thus phasing errors have different effects on species tree estimation depending on whether the species tree is deep or shallow. We suggest that this may be explained by the probability that the sequences from the same species coalesce before they reach the time of species divergence, when one traces the genealogical history at each locus backwards in time. For example, the probability that *S* = 2 sequences from species *A* coalesce before reaching the common ancestor of *A* and *B* is ℙ{*t* mrca *< τ AB*}= 1 *−* e^*−*4^ *≈* 0.982 in the two deep trees and 1*−* e^*−*0.4^ *≈* 0.330 in the two shallow trees, while the corresponding probabilities for *S* = 4 sequences are 0.967 and 0.077 for the deep and shallow trees, respectively (Fig. S4). In the deep trees, there is a high chance for all sequences from the same species to coalesce before reaching species divergence, and then the problem will be similar to using the ancestral sequence for each species (which is mostly determined by the most common nucleotides at the individual sites; Yang *et al*., 1995) for species tree estimation, a process that is not expected to be sensitive to phasing errors. In the shallow species trees, there are high chances that sequences from the same species may not have coalesced before reaching the time of species divergence, and sequences with phasing errors will enter ancestral populations, interfering with species tree estimation.

While our main objective in this study is to evaluate the impacts of different phasing strategies, it is worth noting the effects of other major factors on species tree estimation that are obvious from our results (Figs. 4, S1–S3 and tables 1, S1–S3). By design species tree B is harder to recover than tree U because tree B has four short branches (for clades *C*_10_, *C*_12_, *C*_13_, and *C*_15_) while tree U has only three (for clades *C*_10_, *C*_11_, and *C*_15_) (Fig. 2). Thus tree B is recovered with much lower probability than tree U by all methods in all parameter settings. We note that the individual clades in tree B are recovered with lower probabilities than those in tree U (tables 1, S1–S3). We speculate that this may be due to the fact that the four short branches in tree B are close together (so that 945 trees around them are nearly equally good) while the three short branches in tree U are far apart (so that only 3 *×* 15 = 45 trees around them are nearly equally good). Because of the symmetry in tree B, the probabilities of recovering clades *C*_10_ and *C*_13_ should be equal, as are those for *C*_12_ and *C*_15_. Differences within each pair reflect the random sampling errors due to our use of only 100 replicates. (Note that clades *C*_11_ and *C*_14_ were always recovered in the simulation.)

The mutation rate had a dramatic impact on the precision and accuracy of species tree estimation. At the higher rate (with *θ* = 0.01 vs. 0.001), the credibility set was smaller, its coverage was higher, and the MAP tree matched the true species tree with higher probability. In our species trees, species divergence times (*τ*) are proportional to *θ* . This allows us to compare the two values of *θ*, mimicking the use of conserved or variable regions of the genome for species tree estimation. Our study focuses on closely related species with highly similar sequences, and data simulated at the high rate contain more variable sites and more phylogenetic information.

The number of loci similarly had a huge impact on species tree estimation. With more loci, inference became more precise (with smaller credibility set) and more accurate (with the MAP tree matching the true tree with greater probability). Increasing the number of loci by 10 fold improves performance for all strategies more than increasing the mutation rate by the same factor.

The number of sequences sampled per species had consistent but relatively small effects on species tree estimation. Changing *S* = 2 to 4 improved the probabilities of recovering the true clades in the true species tree, reduced the CI set size, and improved the coverage of the CI set, but the improvements are in general small.

It is noteworthy that the coverage of the 95% CI set was below the nominal 95% in small or uninformative datasets while above 95% in large and informative datasets. In the case of 10 loci at the low rate for tree Deep B, coverage was even below 50% (table S1). Even though the set included nearly 500 trees, more than a half of the CI sets failed to include the true tree. In contrast, at the high mutation rate and with 50 or 100 loci, CI coverage was often 100%. The method is over-confident in small and uninformative datasets and conservative in large and informative ones. The same pattern was noted in a previous simulation examining the information content in phylogenomic datasets (Huang *et al*., 2020, table 3). Note that in our simulation, the replicate datasets are generated under a fixed model (species tree) and fixed parameter values, so that we are evaluating the Frequentist properties of Bayesian model selection, and a match is not expected (Huelsenbeck and Rannala, 2004; Yang and Rannala, 2005). Yet the large discrepancies are striking.

**Table 3.** Mean and standard deviation (*×* 10^*−*3^) of estimates of *θ* for a single population (true value is 0.01) from a sample of *S* sequences using BPP with different strategies of phase resolution and two summary methods

| Method | $S = 2$ | $S = 4$ | $S = 8$ | $S = 16$ | $S = 32$ |
| --- | --- | --- | --- | --- | --- |
| BPP (F) | $10.06 \pm 1.02$ | $10.06 \pm 0.61$ | $10.03 \pm 0.52$ | $9.96 \pm 0.36$ | $10.03 \pm 0.34$ |
| BPP (D) | $10.06 \pm 1.02$ | $10.05 \pm 0.62$ | $10.17 \pm 0.53$ | $10.50 \pm 0.43$ | $11.19 \pm 0.47$ |
| BPP (P) | $10.06 \pm 1.02$ | $9.80 \pm 0.61$ | $9.84 \pm 0.51$ | $9.94 \pm 0.37$ | $10.05 \pm 0.34$ |
| BPP (R) | $10.06 \pm 1.02$ | $12.86 \pm 0.91$ | $18.13 \pm 1.32$ | $26.43 \pm 1.65$ | $41.27 \pm 3.22$ |
| Watterson ($\hat{\theta}_S$) | $9.94 \pm 1.01$ | $9.92 \pm 0.61$ | $9.85 \pm 0.55$ | $9.76 \pm 0.40$ | $9.82 \pm 0.36$ |
| Pairwise distance ($\hat{\theta}_\pi$) | $9.94 \pm 1.01$ | $9.94 \pm 0.63$ | $9.87 \pm 0.63$ | $9.78 \pm 0.50$ | $9.93 \pm 0.55$ |
| Pairwise distance ($\hat{\theta}'_\pi$) | $10.01 \pm 1.03$ | $10.01 \pm 0.64$ | $9.94 \pm 0.64$ | $9.84 \pm 0.50$ | $9.99 \pm 0.56$ |

Note.— Watterson's estimate ( $\hat{\theta}_S$ ) and the average pairwise distance ( $\hat{\theta}_\pi$ ) do not depend on phase resolutions. JC correction is applied in calculation of $\hat{\theta}'_\pi$ .

### Estimation of Divergence Times and Population Sizes under the MSC Model

#### The impact of the phasing strategies

The same datasets simulated for species tree estimation were analyzed to estimate the parameters in the MSC model (*θ*s and *τ*s) with the species tree fixed (Figs. 2a, a′, b & b′). The posterior means and 95% HPD CI for the 100 replicates are plotted in Figures 5, S5–S11, while the relative root mean square errors (rRMSE) are presented in tables S4–S11. Whereas the rRMSE reflects both biases and variances in parameter estimation, the datasets generated by the four phase-resolution strategies have about the same size in terms of the number of loci, the number of sequences per locus, and the number of sites per sequence, so that the sampling errors or variances are similar among methods and the differences in rRMSE mainly reflect differences in bias. Furthermore, we may use the symmetry of species tree B to gauge the magnitude of random sampling errors due to our use of 100 replicates: for instance, rRMSE should be equal for *θ*_*A*_, *θ*_*B*_, *θ*_*E*_ and *θ*_*F*_, and for *τ*_10_ and *τ*_13_, on the balanced trees.

**Fig. 5.**
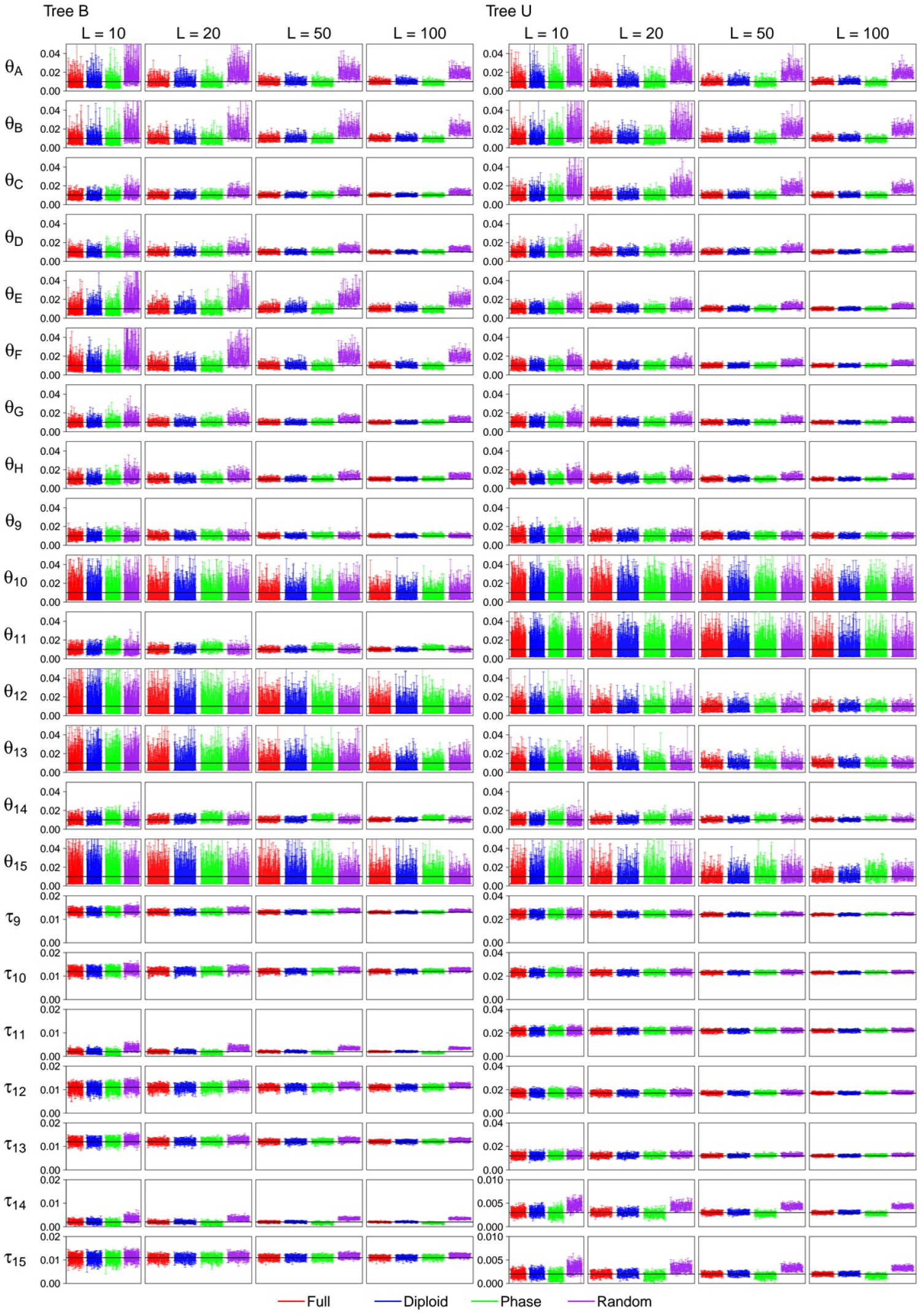
(**MSC, high rate, shallow**, *S* = 4) The 95% HPD CIs for parameters for four phase-resolution strategies: F (the full data), D (diploid), P (PHASE), and R (random) in 100 replicate datasets simulated under MSC model trees Shallow B and Shallow U (Figs. 2a′&b′), at the high mutation rate (*θ* = 0.01) and *S* = 4 sequences per species. The horizontal black lines indicate the true values. Results for other simulation settings are in Figures S5-S11.

The four phase-resolution strategies (F, D, P, and R) performed similarly for the Deep trees at the lower rate and when only *S* = 2 sequences (or one individual) are sampled per species. We note that with *S* = 2 and at the low mutation rate (with heterozygosity at *θ* = 0.001), there will be on average 0.5 heterozygous sites at the same locus, and the probability of having two or more heterozygous sites is 1 *−* 0.999^500^*−* 500 *·* 0.999^499^*·* 0.001 = 0.0901. Then phase resolution will not be a serious issue, and all four strategies examined in the study will be nearly equivalent.

At the high mutation rate (*θ* = 0.01) for the Shallow trees, differences were noted among the strategies even for *S* = 2 sequences (Fig. S6 and tables S5 & S7). The PHASE program produced underestimates for the youngest species divergence times (*τ*_11_ and *τ*_14_ on Shallow B and *τ*_15_ on Shallow U) (Fig. 2a′&b′). The biases became more pronounced when *S* = 4 sequences per species are in the sample (Fig. 5 and tables S9– S11). At the high rate, there are on average 5 heterozygotes per locus in the individual and the probability of having two or more heterozygotes at the locus is 96%. Two factors may be responsible for the bias. First the PHASE program may have inferred heterozygote phase incorrect (indeed the error rate is comparable to that of random phasing with *S* = 2). Second PHASE is an MCMC program generating a distribution of different phase resolutions but we used only the optimal resolution, which may lead to underestimation of sequence divergences.

At the high rate and for shallow trees, random phasing (R) also created serious biases, but the biases are in the opposite direction. Random phasing overestimated the youngest species divergence times (*τ*_11_ and *τ*_14_ on Shallow B and *τ*_15_ on Shallow U), and overestimated *θ* for all modern species. The underestimation of modern *θ* is most striking, and occurred for both deep and shallow species trees at the high rate and is more dramatic with more sequences (*S* = 4 rather than 2) or more loci.

We examined the number of distinct site patterns in the alignment at each locus for the high-rate data (Fig. S12). Site patterns are compressed for the JC model, so that one site pattern is constant while the others are variable (Yang, 2006, p.144), and the number is thus an indication of the level of sequence divergence. At almost every locus, the PHASE program (P) produced alignments with fewer distinct site patterns than the true phase resolution (for example, with the mean to be 36.07 compared with the true value 38.51 on tree B), apparently because we used the optimal phase resolution inferred by the program and ignored the less likely ones. Random resolution produced about the same number of site patterns as the true number (average 38.36 vs. 38.51 for tree Deep B). The number of site patterns is thus not the reason for the poor performance of random phasing.

Note that calculation of the heterozygosity for each diploid individual, which is simply the proportion of heterozygous sites in the two sequences at the locus, does not rely on phase resolution. If we calculate the heterozygosity for each diploid individual and then average over individuals of the same species, we will get a reasonably good estimate of *θ* for that species. However, in the gene-tree based analysis conducted in BPP, each randomly phased haploid sequence is compared not only with the other sequence from the same individual, but also with sequences from other individuals through the use of a gene tree relating all phased haploid sequences at the locus. While the true haploid sequences may all be closely related, random phase resolution may generate chimeric sequences that are very different from naturally occurring fully resolved sequences, inflating apparent coalescent times and genetic diversity in the population. This effect is expected to be more serious when more individuals are included in the sample.

#### Estimation of *θ* for a single species

To explore this interpretation, we conducted a small simulation sampling independent loci from a single species to estimate the only parameter *θ* (Fig. 6, table 3). With *S* = 2 sequences per locus (one diploid individual), the four phase-resolution strategies are equivalent. However, with the increase of *S*, the strategy of random phase resolution becomes increasingly biased. Previously Felsenstein (1992) examined the efficiency of two summary methods based on the number of segregating (variable) sites (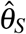; Watterson, 1975) and the average pairwise distance (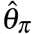 ; Tajima, 1983), relative to the maximum likelihood (ML) method based on gene genealogies. He found that the summary methods (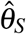 and 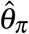) were much less efficient than the ML estimate, with orders-of-magnitude differences in the variance in large samples (Felsenstein, 1992, tables 1 and 2), indicating that there is much information about *θ* in the genealogical histories. The ML method should be very similar to BPP here as both are full likelihood methods. Here we note that the number of segregating sites does not depend on phase resolutions, and similarly the average proportion of different sites, averaged over all the *S*(*S −* 1)*/*2 pairwise comparisons, depends on the site configurations at each variable site (such as 10 Ts and 4 Cs) but not on the genotypic phase between different heterozygous sites. Both Waterson’s estimator and the average pairwise distance are thus unaffected by phasing errors. It is also noteworthy that those two simple methods are not affected by recombination within the locus, while coalescent-based methods are (Felsenstein, 2019). While it is not unexpected that a full likelihood method may be more sensitive to certain errors in the model or in the data than heuristic methods, in this case it is striking that the systematic bias is so large (with estimates to be several times larger than the true value) when the coalescent-based method is applied to randomly phased sequences.

**Fig. 6.**
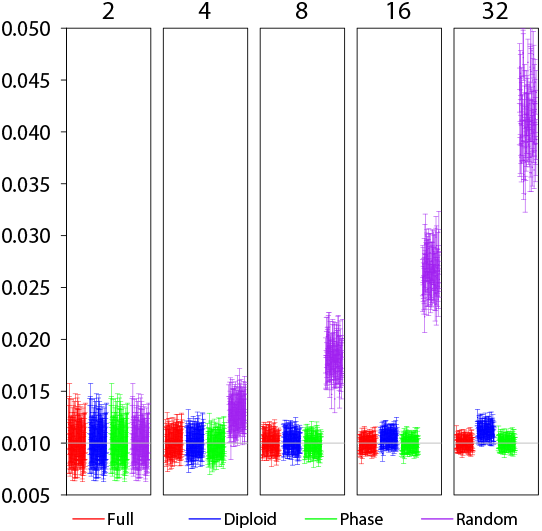
The 95% HPD CIs for parameter *θ* in the single-population coalescent model in 100 replicate datasets using four phase-resolution strategies: F (the full data), D (diploid), P (PHASE), and R (random). There are 100 independent loci in each dataset, and at each locus there are *S* sequences of 500 sites (or *S/*2 diploid individuals), with *S* = 2, 4, 8, 16, and 32. The true parameter value is 0.01.

Felsenstein’s (1992) analysis, as mentioned above, assumed knowledge of the true gene trees and coalescent times (or equivalently infinitely long sequences at each locus). Here BPP is applied to sequence alignments and accommodates uncertainties in the genealogical trees. The different methods then have much more similar performance (table 3, 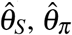 and BPP strategy F), suggesting that the uncertainties in the genealogical trees due to mutational variations in the sequences have eroded much of the information in the gene trees. The summary methods (in particular, 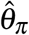) have larger variances than the BPP estimates, especially in large samples of *S* = 32 sequences, but the differences are relatively small. We also note that analytical phase integration (D) produced variances that are nearly identical to those for the use of the full data (F).

#### Impacts of other factors on parameter estimation under the MSC model

We note that different parameters are estimated with very different precision and accuracy, reflecting the different amount of information in the data. Population size parameters (*θ*s) for modern species are well estimated, as well as *θ*_9_ for the root population, but *θ*s for other ancestral species, especially those represented by very short branches (e.g., *θ*_10_, *θ*_13_, *θ*_12_, *θ*_15_ in tree B) have large errors (Figs. 5, S5–S11). Species divergence times are all well estimated, with rRMSE to be even much smaller than those for population size parameters for modern species (tables S4–S11).

Both the mutation rate and the number of loci had a major impact on the estimation of the parameters. For all phasing strategies increasing the number of loci by 10 fold improves performance more than increasing the mutation rate by the same factor (Figs. 5, S5–S11, tables S4–S11).

### Estimation of Introgression Probability under the MSci Model

We used the MSci models of Figure 2c&c′ to simulate sequence data and used BPP to analyze them to estimate parameters in the MSci model. We are in particular interested in whether the different strategies of heterozygote phase resolution may lead to biases in the estimation of the timing (*τ*_*H*_) and strength of the introgression (*ϕ*). The results are summarized in Figures 7 & S13–S19 and tables 4 & S12–S18.

**Table 4.** (**MSci A00** *S* = 4, **high rate, shallow)** Relative root mean square error (rRMSE) for parameter estimates under the Deep MSci model (fig. 2c′) with *ϕ* = 0.1 or 0.3 at the high mutation rate and *S* = 4

| Truth | $\theta_A$ | $\theta_B$ | $\theta_C$ | $\theta_R$ | $\theta_S$ | $\theta_T$ | $\theta_H$ | $\tau_R$ | $\tau_S$ | $\tau_T$ | $\phi$ | |
| --- | --- | --- | --- | --- | --- | --- | --- | --- | --- | --- | --- | --- |
|  | 1 | 1 | 1 | 1 | 1 | 1 | 1 | 3 | 1 | 2 | 0.1 | 0.3 |
| $\phi = 0.1$ | | | | | | | | | | | | |
| F, 10L | 0.28 | 0.24 | 0.34 | 0.24 | 0.45 | 0.36 | 0.25 | 0.15 | 0.45 | 0.17 | 0.42 | - |
| D, 10L | 0.31 | 0.25 | 0.36 | 0.24 | 0.48 | 0.36 | 0.22 | 0.16 | 0.46 | 0.18 | 0.41 | - |
| P, 10L | 0.27 | 0.22 | 0.28 | 0.27 | 0.50 | 0.51 | 0.27 | 0.15 | 0.45 | 0.22 | 0.43 | - |
| R, 10L | 0.99 | 0.86 | 1.49 | 0.31 | 0.37 | 0.25 | 0.23 | 0.36 | 1.12 | 0.46 | 0.47 | - |
| F, 20L | 0.24 | 0.21 | 0.28 | 0.18 | 0.40 | 0.35 | 0.28 | 0.10 | 0.41 | 0.13 | 0.46 | - |
| D, 20L | 0.26 | 0.22 | 0.31 | 0.18 | 0.43 | 0.33 | 0.26 | 0.11 | 0.43 | 0.14 | 0.45 | - |
| P, 20L | 0.24 | 0.20 | 0.26 | 0.26 | 0.39 | 0.49 | 0.38 | 0.11 | 0.46 | 0.17 | 0.54 | - |
| R, 20L | 1.09 | 0.73 | 1.75 | 0.27 | 0.32 | 0.25 | 0.22 | 0.30 | 1.04 | 0.41 | 0.53 | - |
| F, 50L | 0.17 | 0.12 | 0.18 | 0.13 | 0.24 | 0.27 | 0.39 | 0.07 | 0.33 | 0.08 | 0.53 | - |
| D, 50L | 0.18 | 0.13 | 0.20 | 0.13 | 0.27 | 0.27 | 0.32 | 0.07 | 0.38 | 0.09 | 0.51 | - |
| P, 50L | 0.31 | 0.13 | 0.34 | 0.22 | 0.21 | 0.41 | 0.29 | 0.09 | 0.60 | 0.14 | 0.68 | - |
| R, 50L | 1.18 | 0.63 | 1.50 | 0.22 | 0.23 | 0.26 | 0.24 | 0.26 | 0.90 | 0.38 | 0.64 | - |
| F, 100L | 0.12 | 0.08 | 0.13 | 0.09 | 0.18 | 0.23 | 0.32 | 0.04 | 0.27 | 0.06 | 0.59 | - |
| D, 100L | 0.14 | 0.08 | 0.16 | 0.09 | 0.19 | 0.24 | 0.30 | 0.04 | 0.33 | 0.06 | 0.57 | - |
| P, 100L | 0.39 | 0.11 | 0.42 | 0.21 | 0.15 | 0.42 | 0.23 | 0.09 | 0.72 | 0.14 | 0.74 | - |
| R, 100L | 1.27 | 0.59 | 1.60 | 0.19 | 0.20 | 0.21 | 0.23 | 0.23 | 0.73 | 0.35 | 0.73 | - |
| $\phi = 0.1$ | | | | | | | | | | | | |
| F, 10L | 0.30 | 0.27 | 0.31 | 0.21 | 0.35 | 0.35 | 0.30 | 0.15 | 0.34 | 0.16 | - | 0.48 |
| D, 10L | 0.31 | 0.29 | 0.41 | 0.22 | 0.40 | 0.35 | 0.28 | 0.16 | 0.35 | 0.17 | - | 0.49 |
| P, 10L | 0.27 | 0.24 | 0.27 | 0.22 | 0.48 | 0.55 | 0.35 | 0.15 | 0.48 | 0.21 | - | 0.44 |
| R, 10L | 1.03 | 0.93 | 1.83 | 0.32 | 0.37 | 0.28 | 0.24 | 0.36 | 1.11 | 0.49 | - | 0.57 |
| F, 20L | 0.24 | 0.18 | 0.26 | 0.18 | 0.35 | 0.28 | 0.37 | 0.09 | 0.30 | 0.12 | - | 0.39 |
| D, 20L | 0.28 | 0.20 | 0.33 | 0.19 | 0.33 | 0.27 | 0.33 | 0.10 | 0.33 | 0.13 | - | 0.40 |
| P, 20L | 0.26 | 0.17 | 0.29 | 0.21 | 0.45 | 0.47 | 0.42 | 0.10 | 0.52 | 0.18 | - | 0.37 |
| R, 20L | 1.04 | 0.69 | 1.69 | 0.30 | 0.30 | 0.26 | 0.24 | 0.31 | 1.08 | 0.44 | - | 0.51 |
| F, 50L | 0.18 | 0.12 | 0.16 | 0.11 | 0.22 | 0.28 | 0.48 | 0.05 | 0.21 | 0.10 | - | 0.27 |
| D, 50L | 0.20 | 0.12 | 0.19 | 0.11 | 0.21 | 0.29 | 0.49 | 0.06 | 0.25 | 0.10 | - | 0.30 |
| P, 50L | 0.28 | 0.13 | 0.34 | 0.20 | 0.26 | 0.53 | 0.56 | 0.09 | 0.60 | 0.17 | - | 0.33 |
| R, 50L | 0.97 | 0.61 | 1.56 | 0.24 | 0.21 | 0.24 | 0.25 | 0.29 | 1.07 | 0.41 | - | 0.38 |
| F, 100L | 0.11 | 0.09 | 0.11 | 0.08 | 0.15 | 0.20 | 0.42 | 0.04 | 0.15 | 0.08 | - | 0.20 |
| D, 100L | 0.12 | 0.10 | 0.14 | 0.08 | 0.14 | 0.20 | 0.49 | 0.04 | 0.19 | 0.08 | - | 0.21 |
| P, 100L | 0.34 | 0.12 | 0.41 | 0.20 | 0.19 | 0.47 | 0.37 | 0.09 | 0.66 | 0.15 | - | 0.34 |
| R, 100L | 0.87 | 0.59 | 1.50 | 0.22 | 0.18 | 0.17 | 0.30 | 0.27 | 1.06 | 0.39 | - | 0.32 |

Note.— Truth represents the true parameter values used in the simulation; values for $\theta$ and $\tau$ are $\times 10^{-2}$ . Results for other simulation settings are in tables S12-S18.

**Fig. 7.**
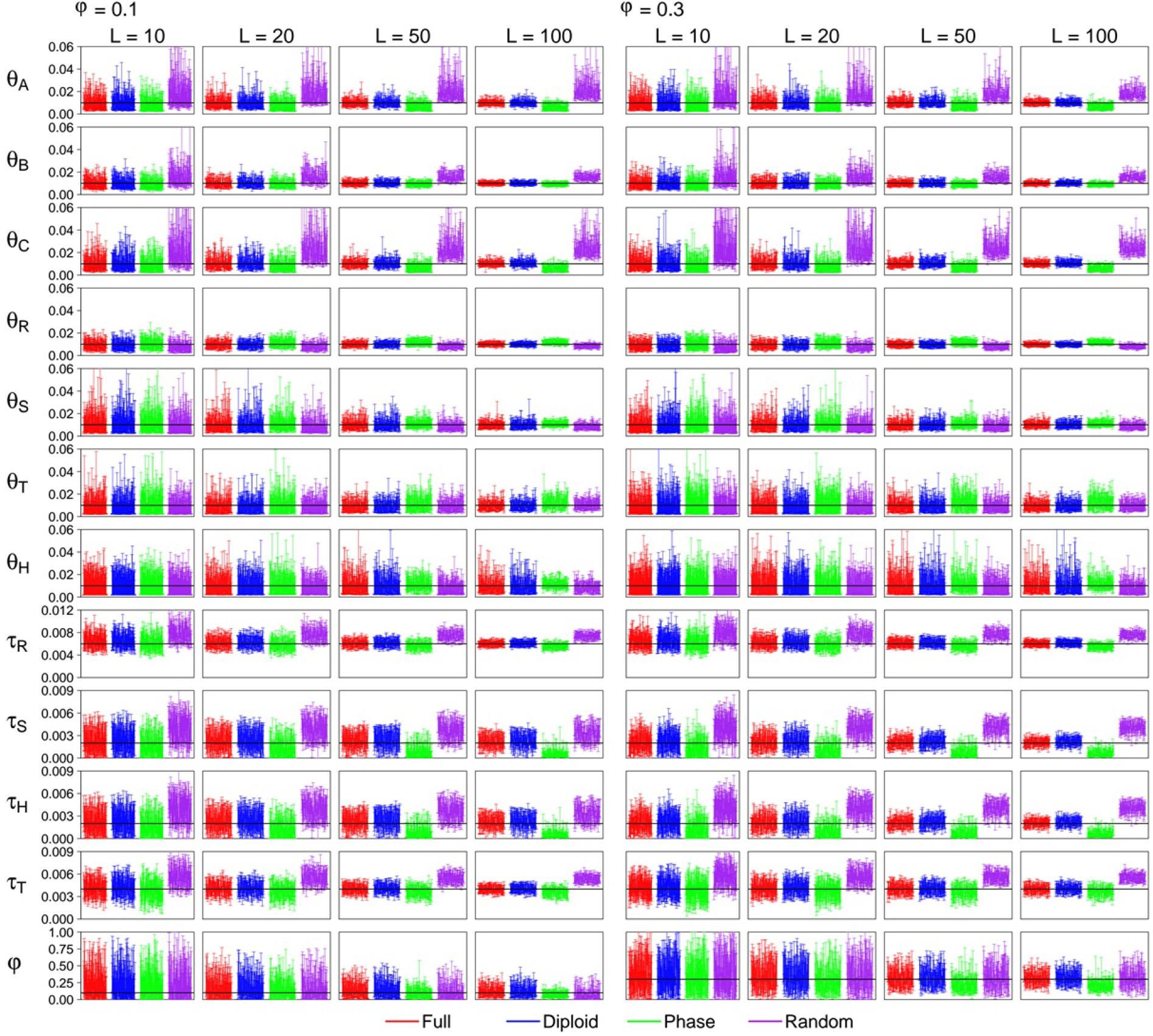
(**MSci model, high rate, Shallow**, *S* = 4) The 95% HPD CIs for parameters under the MSci model of Figure 2c′ when *S* = 4 sequences are sampled per species. Results for *S* = 2 are in Figure S14. See legend to Figure 5.

As before, the diploid strategy (D) produced results almost indistinguishable from the use of the full data (F) in all parameter settings. The performance of the PHASE program (P) and random phasing (R) depends on the mutation rate and, to an lesser extent, on the number of sequences per species *S*. At the low rate, and in particular with only *S* = 2 sequences per species, all four strategies have similar performance, but large differences were found at the high mutation rate. Strategy R overestimates the modern *θ* and the species divergence times (*τ*) at the high rate, with the bias being more serious for *S* = 4 sequences than for *S* = 2. This is the same behavior as discussed earlier in the simulation under the MSC model. Strategy R also tends to overestimate *φ*, but the bias is small. Strategy P had the opposite bias and produced underestimates of modern *θ* and species divergence times when the mutation rate is high, with smaller biases than for strategy R. Strategy P also underestimates the introgression probability (*φ*).

An interesting question is whether each method detects introgression. We calculated the proportion of replicates in which the lower limit of the 95% HPD CI for *φ* exceeds a small value, set somewhat arbitrarily at 0.001. If the CI excludes the small value, we may take it as evidence that *φ* = 0 is ruled out so that there is significant evidence for introgression. By this measure of power of the Bayesian ‘test’, strategies D and P had nearly identical power as the use of the full data (F), while random resolution (R) had reduced power at the high mutation rate (tables 5 & S19). Overall, power was very high even with only 10-20 loci and at the low mutation rate. Having more sequences is noted to boost the power of the test for all phase-resolution strategies.

**Table 5.** (**MSci test, shallow)** Power of the Bayesian test for introgression (measured by the proportion of replicates in which the lower limit of the 95% HPD CI for *φ* is *>* 0.001) when the true model is the Shallow MSci tree

|  | low |  |  |  | high |  |  |  |
| --- | --- | --- | --- | --- | --- | --- | --- | --- |
|  | 10L | 20L | 50L | 100L | 10L | 20L | 50L | 100L |
| $\phi = 0.1$ | | | | | | | | |
| $S = 2$ seqs per species | | | | | | | | |
| F | 0.42 | 0.41 | 0.42 | 0.49 | 0.48 | 0.52 | 0.64 | 0.89 |
| D | 0.48 | 0.40 | 0.45 | 0.46 | 0.38 | 0.35 | 0.61 | 0.80 |
| P | 0.49 | 0.45 | 0.49 | 0.49 | 0.55 | 0.60 | 0.83 | 0.99 |
| R | 0.51 | 0.44 | 0.47 | 0.48 | 0.27 | 0.25 | 0.40 | 0.57 |
| $S = 4$ seqs per species | | | | | | | | |
| F | 0.58 | 0.47 | 0.55 | 0.58 | 0.59 | 0.71 | 0.98 | 0.99 |
| D | 0.56 | 0.44 | 0.54 | 0.60 | 0.56 | 0.66 | 0.94 | 0.99 |
| P | 0.57 | 0.43 | 0.44 | 0.49 | 0.60 | 0.65 | 0.94 | 1.00 |
| R | 0.56 | 0.45 | 0.50 | 0.57 | 0.44 | 0.46 | 0.71 | 0.81 |
| $\phi = 0.3$ | | | | | | | | |
| $S = 2$ seqs per species | | | | | | | | |
| F | 0.68 | 0.74 | 0.87 | 0.94 | 0.86 | 0.97 | 1.00 | 1.00 |
| D | 0.64 | 0.81 | 0.89 | 0.93 | 0.84 | 0.91 | 1.00 | 1.00 |
| P | 0.68 | 0.78 | 0.85 | 0.89 | 0.86 | 0.94 | 0.99 | 1.00 |
| R | 0.66 | 0.72 | 0.87 | 0.92 | 0.74 | 0.82 | 0.96 | 0.99 |
| $S = 4$ seqs per species | | | | | | | | |
| F | 0.79 | 0.88 | 0.97 | 1.00 | 0.95 | 1.00 | 1.00 | 1.00 |
| D | 0.83 | 0.86 | 0.95 | 1.00 | 0.92 | 0.99 | 1.00 | 1.00 |
| P | 0.84 | 0.84 | 0.95 | 1.00 | 0.94 | 0.97 | 1.00 | 1.00 |
| R | 0.84 | 0.82 | 0.94 | 0.99 | 0.75 | 0.89 | 1.00 | 1.00 |

Note.— Results for the Deep MSci model are in table S19.

**Table 6.**
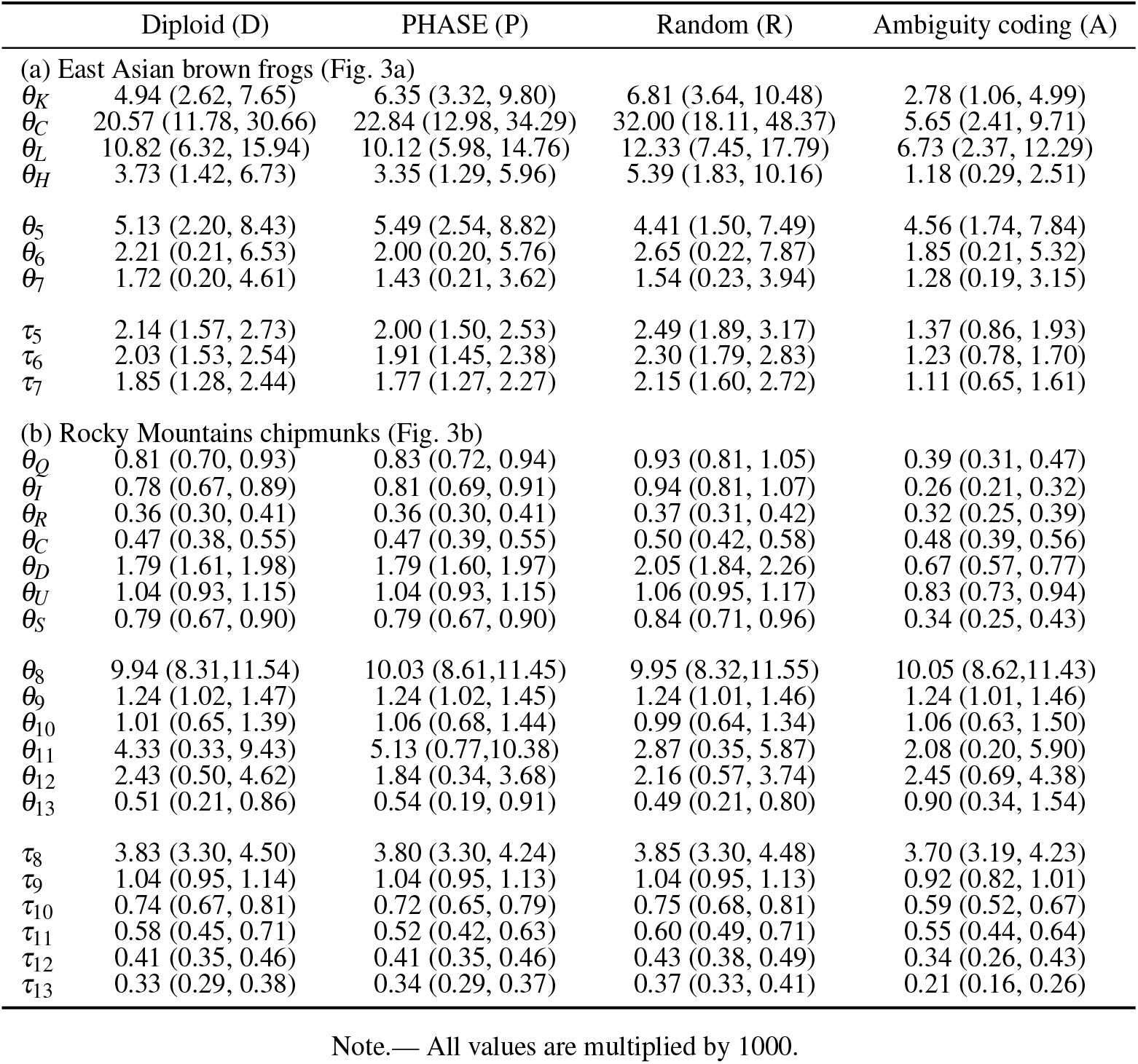
Posterior means and 95% HPD CIs for parameters under the MSC model for the east Asian brown frogs and for the chipmunks

### Running Time for Different Analyses

The running time for the A01 analysis under the MSC model (species tree estimation) for the four phasing strategies (F, D, P, and R), averaged over the 100 replicates, is plotted against the number of loci in Figure S20. Running time increases nearly linearly with the number of loci, with the slope being steeper when *S* = 4 sequences are sampled per species than for *S* = 2. The diploid integration algorithm (D) has the longest running time. Note that the number of parameters in the MSC model, the number of loci, the number of sequences etc. are identical for the four strategies, so that their computational load is proportional to the number of site patterns. As strategy D enumerates all possible phase resolutions (including the true resolution), which may result in many distinct site patterns, it is more expensive than the other methods. The running time for each BPP analysis on a single core ranged from *∼* 20 minutes for 10 loci to *∼* 5 hours for strategy D with data of 100 loci. Strategy P involves running the Bayesian MCMC program PHASE for each of the *L* loci. At the low mutation rate with very few heterozygous sites per locus, this requires minimal computation (Fig. S21), but at the high rate and with *S* = 4 sequences per species, the running time can be comparable with running the subsequent BPP analyses.

The running time for the A00 analysis (parameter estimation) under the MSC and MSci models is shown in Figures S22–S25. The A00 analysis under the MSC involves less computation than the A01 analysis as there is no MCMC moves to change the species tree. Overall, the same patterns are observed as discussed above for the A01 analysis. Note that the computer cluster used in this work consist of computers with different processors, so there may be random fluctuations in running time due to the different jobs being assigned to different processors. For example the differences in Figures S21 & S23 reflect this random fluctuation as the data were the same.

### Analysis of two real datasets

We analyzed two real datasets using four different phase-resolution strategies: D (diploid), P (PHASE), R (random), and A (ambiguity). With real data, the option of true phase resolution (F) is unavailable, and the analytical phase resolution (D) is expected to have the best performance, against which we compare the other strategies.

#### East Asia brown frogs

We re-analyzed a dataset of five nuclear loci from the East Asia brown frogs in the *Rana chensinensis* species complex (Zhou *et al*., 2012) (Fig. 3a). This dataset was previously analyzed by Yang (2015) using strategy A. The number of site patterns at each locus is 18–26 for strategy A, and 22–102 for strategy D. Running time using one thread on our server was 3 mins for A, 7-8 mins for P and R, and 12 mins for D.

In the A01 analysis (species tree estimation), the four strategies (D, P, R, and A) produced the same MAP tree (Fig. 3a): (((H, L), C), K), with the posterior to be 0.29 for D, 0.36 for P, 0.35 for R, and 0.21 for A. The analysis of Yang (2015) produced a different MAP tree, ((H, L), (C, K)). The difference is due to the use of different priors: Yang (2015) used BPP3.1, with gamma priors on the parameters (*θ*s for all populations and *τ* for the root), whereas here inverse gamma priors are used in BPP4.3. Note that the species trees have low support in both analyses.

In the A00 analysis (parameter estimation under MSC with the MAP species tree fixed), the posterior means and 95% HPD intervals are shown in table 6a. Strategy P (PHASE) produced similar results to strategy D. Strategy R (random) produced overestimates of *θ*s for modern species, while strategy A (ambiguity) produced serious underestimates of *θ*s for modern species and divergence times. The results are consistent with our findings from the simulation.

#### Rocky Mountains chipmunks

In the A01 analysis (species tree inference) of the 500 nuclear loci for Rocky Mountains chipmunks, strategies D, P, and R produced the same MAP tree, shown in Figure 3b, with the posterior for every node *∼* 1.0. This is also the species tree inferred by Sarver *et al*. (2021) using summary methods, although the authors obtained lower support values even with all 1060 loci used. The difference may be due to the higher power of the BPP analysis, which uses the full data rather than data summaries (e.g., Shi and Yang 2018; Kim and Degnan, 2020; Zhu and Yang, 2021). Strategy A (ambiguity) produced a different MAP species tree from the other strategies (Fig. 3b), with the relationship (C, (D, (IQR))) instead of (D, (C, (IQR))), with the posterior at 0.94. The running time for the A01 analysis, using eight cores on a server with Intel Xeon Gold 6154 3.0GHz processors, was 9 hours for strategy A, and 16-17 hours for strategies D, P, and R, with strategy D having slightly longer running time. The number of site patterns at the 1060 loci for strategy D is shown in Fig. S26. Strategy P also needed the additional time for running the PHASE program, which was 33 mins to phase all 1060 loci using one thread on the server.

In the A00 analysis (parameter estimation), strategy P (PHASE) produced nearly identical results to strategy D (diploid) (table 6). Compared with strategy D, strategy R (random) produced overestimates of *θ*s for modern species, while divergence times for recent nodes were also over-estimated very slightly. Strategy A (ambiguity) produced serious underestimates of *θ*s for modern species, with divergence times, especially of recent nodes, to be underestimated as well. Those results mimic our findings about the relative performance of the different strategies in the simulated data. Running time for the A00 analysis was 2.5 hours for strategy A, and 5-6 hours for strategies D, P, and R. Note that in the A00 analysis the chain is only half as long as in the A01 analysis.

## 4. Discussion

### The Impact of Phasing Errors Depends on the Inference Problem

We have used simulation to examine the performance of four different strategies for handling heterozygote phase in genomic sequence data: F (full phased data), D (diploid analytical phase integration), P (PHASE), and R (random). Inference problems examined have included species tree estimation under the MSC model and parameter estimation under the MSC and MSci models. We found that the different strategies, including random phase resolution (or equivalently the use of haploid consensus sequences), did not affect species tree estimation when the species divergences are much older than the coalescent times. The different phasing strategies may be expected to have even less impact on inference of deep phylogenies, where within-species polymorphism is much lower than between-species divergence. However, species tree estimation is affected by phasing errors if the species tree is shallow and between-species divergence is similar to within-species polymorphism, if the mutation rate is high so that there are many heterozygote sites in the sequence, and if many sequences are sampled from each species. Phasing errors are clearly important when genomic data are used to infer the divergence history of populations of the same species.

We found that estimation of parameters in the MSC and MSci models is more sensitive to phasing errors than is species tree estimation. In particular, population sizes for modern species are seriously overestimated under the MSC and MSci models when random phasing or haploid consensus sequences are used. Our analysis of the simple case of estimating *θ* under the single-population coalescent suggests that the bias is caused mainly by the unusual sequences generated by random phase resolution (Fig. 6 and table 3). Estimates of the introgression probability and introgression time under the MSci model may also be biased by errors in random phasing. The biases are more serious when the mutation rate is high so that there are multiple heterozygote sites at each locus and when multiple sequences are sampled per species. Those results are consistent with Gronau *et al*. (2011), who also found that random phase resolution affected parameter estimation in their analysis of genomic sequence data from different human populations.

### Limitations of our Simulation and Implications to Practical Data Analysis

Here we note a few limitations of our study. First we have examined only one inference method, the Bayesian method implemented in the BPP program. Our results may be expected to apply to other full likelihood implementations such as StarBeast (Ogilvie *et al*., 2017; Zhang *et al*., 2018) or PhyloNet-Seq (Wen and Nakhleh, 2018), but may not apply to summary methods. Similarly we considered only a few inference problems under the MSC and MSci models using genomic sequence data. We have not examined the impact of phasing errors on inference of population demographic changes or on inference of migration/introgression histories (our simulation under the MSci model assumed a fixed introgression event).

Given those caveats, we discuss the implications of our simulation results to practical data analysis. First, our simulation as well as those of Gronau *et al*. (2011) and Andermann *et al*. (2019) suggest that random phase resolution or the use of haploid consensus sequences should be avoided. Strategy R never performed better than computational phasing (strategy P) in our simulations. Similarly strategy A (ambiguity) should not be recommended. Virtually all phylogenetic likelihood programs accommodate ambiguities in a sequence alignment representing undetermined nucleotides using a data augmentation algorithm in the likelihood calculation (Felsenstein, 2004, pp.255–6; Yang, 2014, pp.110-112). As heterozygotes (with, e.g., Y meaning both T and C) are not ambiguities (with Y meaning either T or C), this approach misinterprets the data, and has the obvious effect of underestimating the heterozygosity or *θ* for the modern species. Bias may also be introduced into estimates of other parameters, such as underestimation of divergence times (Andermann *et al*., 2019). The approach also underestimates the information content in the data, as it in effect treats two sequences (although unphased) as only one. This mistake in the treatment of the data was made by Rannala and Yang (2003) in the analysis of three human noncoding loci of Zhao *et al*. (2000), Yu *et al*. (2001), and Makova *et al*. (2001), and by Yang (2015) in the analysis of the five nuclear loci from East Asian brown frogs (Zhou *et al*., 2012). The mistake is easy to see from the occurrence of the same ambiguity character (such as Y) in multiple sequences at the same site in the alignment.

Strategy D (diploid analytical integration) produced results that are extremely similar to the use of the full data (F) in all simulation settings of this study (see also Gronau *et al*., 2011). As the algorithm averages over all possible phase resolutions and constitutes a full likelihood approach to handling missing data, it is the optimal statistical approach when the data consist of unphased diploid sequences, and may thus be recommended in general, even for inference problems that are not examined in our simulation study. As a statistical inference method, strategy D is equivalent to the approach of sampling phase resolutions in a Markov chain Monte Carlo (MCMC) algorithm (Kuhner and Felsenstein, 2000). In small or intermediate datasets, analytical phase integration appears more efficient computationally than MCMC, whereas for large datasets, both may be unfeasible.

Note that analyses under the four strategies F, D, P, and R involve the same number of species, the same number of parameters, the same number of loci, the same number of sequences, etc., with the only difference being in the number of site patterns. The relative computational load for the strategies is thus proportional to the number of site patterns. Strategy D performs phylogenetic likelihood calculation (Felsenstein, 1981) for all distinct site patterns that may result from enumerating all possible phase resolutions, which include the true phase resolution. Thus strategy D involves at least as many site patterns as in the full data (strategy F). For the simulations of this study, the number of site patterns for strategy D is less than twice the number for strategy F (Fig. S12). However, if there are many long sequences of high heterozygosity at a locus, enumeration of all phase resolutions may lead to a huge number of site patterns. For example, the three noncoding regions of human DNA analyzed by Rannala and Yang (2003) have about 60 sequences per locus, with *∼* 10^4^ sites. The number of site patterns in the unphased alignments (strategy A) is 50–73, but reaches 1.2–4.4 million for strategy D, rendering the analysis unfeasible. Note that those loci are long genomic segments, which may be affected by recombination, whereas datasets suitable for analysis under the MSC typically involve much shorter genomic segments (e.g., Burgess and Yang, 2008).

We suggest that computational phasing (strategy P) should be an acceptable alternative when strategy D is computationally unfeasible. In our analyses of the simulated and real datasets, strategy P produced similar results to the use of full data (F) or the analytical phase integration approach (D), with very small biases. Note that the Bayesian program PHASE assumes a population genetics model and is designed for sequence or allelic data from the same species.

However, our use of it to analyze sequence data from multiple species produced relatively small biases in parameter estimation in both simulated data and in the two real datasets, much better than random phase resolution or haploid consensus sequences. We also note that phasing based on reads combined with bioinformatic analysis shows great promise (Andermann *et al*., 2019). In particular, exciting developments in sequencing technology to provide longer reads, combined with computational algorithms (Porubsky *et al*., 2020; Zhou *et al*., 2020; Cheng *et al*., 2021), may soon make it practical to produce routinely fully phased diploid genomes.

## 5. Supplementary Material

Data available from the Dryad Digital Repository: https://doi.org/10.5061/dryad.vmcvdncrd.

## 6. Acknowledgments

We thank two anonymous reviewers, the AE (Laura Kubatko) and the EiC (Bryan Carstens) for constructive comments. We thank Jiayi Ji for converting the chipmunk alignments of Sarver *et al*. (2021) from the Nexus format to the BPP format.

## 8. Supplementary Material

Data available from the Dryad Digital Repository: http://dx.doi.org/10.5061/dryad.xxxxxx.

